## Supplemental file 4 for "Microbe transmission from pet shop to lab-reared zebrafish reveals a pathogenic birnavirus"

A

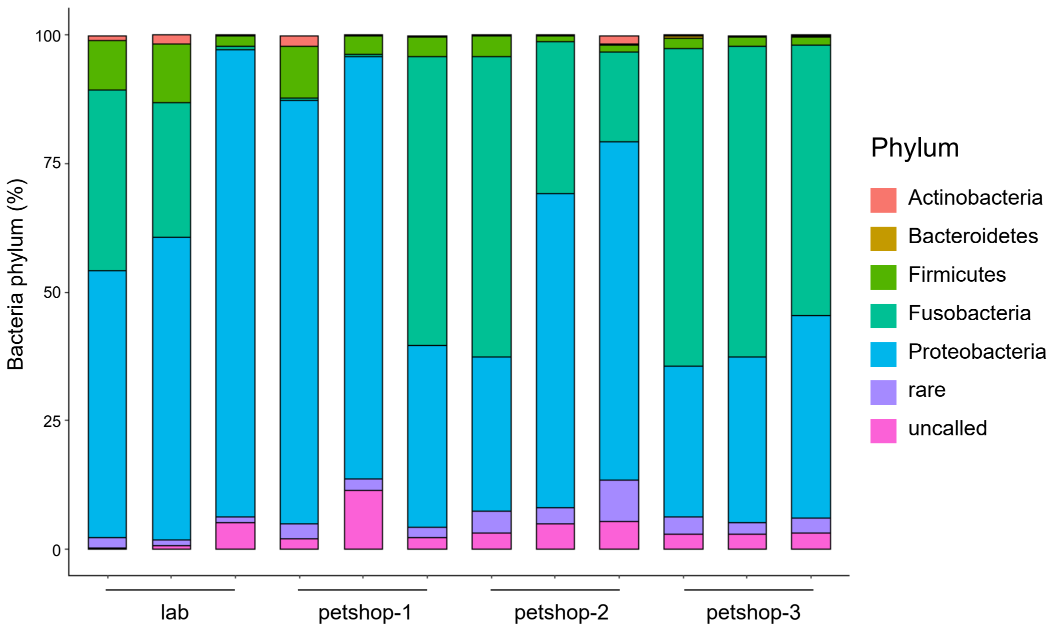


B

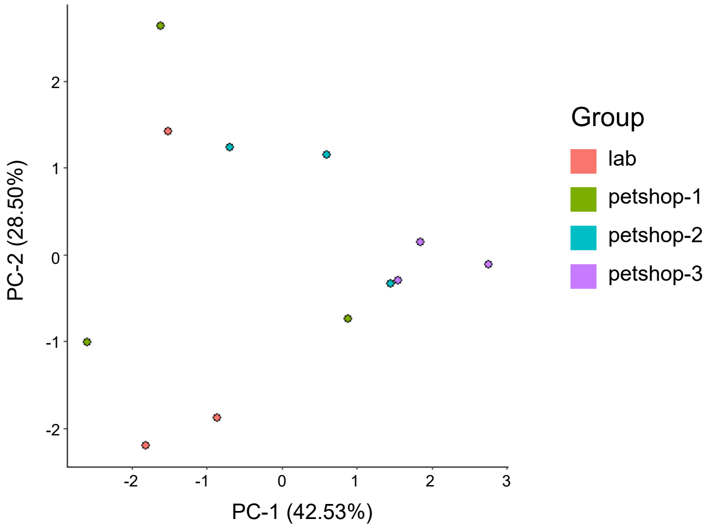


**Supplemental file 4 Gut microbiome composition of lab and pet store zebrafish.** A) bar graphs illustrating bacteria relative abundance by phylum. Relative abundance was calculated form normalized read counts from each sample. Each column represents an individual fish intestine sample. N = 3 for each group. Note large degree of variation between individuals and between groups. B) Principal component analysis using relative abundances of each bacteria phylum per individual zebrafish. PC-1 and PC-2 combined account for ~ 70% of variation between samples. Note large variation within and between groups. Small sample size per group (n = 3) prevented us from plotting a 95% confidence ellipse for each group. A & B) generated with ggplot2 in R studio.
